# Computational workflow and interactive analysis of single-cell expression profiling of islets generated by the Human Pancreas Analysis Program

**DOI:** 10.1101/2023.01.03.522578

**Authors:** Abhijeet R. Patil, Jonathan Schug, the HPAP Consortium, Ali Naji, Klaus H. Kaestner, Robert B. Faryabi, Golnaz Vahedi

## Abstract

Type 1 and Type 2 diabetes are distinct genetic diseases of the pancreas which are defined by the abnormal level of blood glucose. Understanding the initial molecular perturbations that occur during the pathogenesis of diabetes is of critical importance in understanding these disorders. The inability to biopsy the human pancreas of living donors hampers insights into early detection, as the majority of diabetes studies have been performed on peripheral leukocytes from the blood, which is not the site of pathogenesis. Therefore, efforts have been made by various teams including the Human Pancreas Analysis Program (HPAP) to collect pancreatic tissues from deceased organ donors with different clinical phenotypes. HPAP is designed to define the molecular pathogenesis of islet dysfunction by generating detailed datasets of functional, cellular, and molecular information in pancreatic tissues of clinically well-defined organ donors with Type 1 and Type 2 diabetes. Moreover, data generated by HPAP continously become available through a centralized database, PANC-DB, thus enabling the diabetes research community to access these multi-dimensional data prepublication. Here, we present the computational workflow for single-cell RNA-seq data analysis of 258,379 high-quality cells from the pancreatic islets of 67 human donors generated by HPAP, the largest existing scRNA-seq dataset of human pancreatic tissues. We report various computational steps including preprocessing, doublet removal, clustering and cell type annotation across single-cell RNA-seq data from islets of four distintct classes of organ donors, i.e. non-diabetic control, autoantibody positive but normoglycemic, Type 1 diabetic, and Type 2 diabetic individuals. Moreover, we present an interactive tool, called CellxGene developed by the Chan Zuckerberg initiative, to navigate these high-dimensional datasets. Our data and interactive tools provide a reliable reference for singlecell pancreatic islet biology studies, especially diabetes-related conditions.

## Background & Summary

The pancreas is part of the gastrointestinal system and is anatomically divided into four parts: head, neck, body, and tail^1^. While most of the pancreas functions as an exocrine accessory digestive gland secreting digestive enzymes into the gut lumen, a small minority of endocrine cells are organized into cell aggregates termed “islets of Langerhans”, or simply “islets”. The islets of Langerhans contain five endocrine cell types, the glucagon (GCG) producing alpha cells, insulin (INS) producing beta cells, ghrelin (GHRL) producing epsilon cells, somatostatin (SST) producing delta cells, and pancreatic polypeptide (PPY) cells^2^. The exocrine pancreas makes up about 80% pancreas volume^3^ and consists of acinar cells producing digestive enzymes and ductal cells that form a conduit for the exocrine secrtions to reach the duodenum^4^. Finally, there are mesenchymal stellate cells which are found both in endocrine and exocrine glands^5^. Previously, multiple researchers have performed transcriptional profiling experiments using microarray and bulk RNA sequencing on pancreatic tissues^6–11^. However, these experiments provide only the average RNA expression and do not reflect the expression at the single-cell level.

A revolution in single-cell transcriptomic technologies of the past decade has also influenced the molecular profiling of islets. Several primary single-cell atlases for murine and humans^12–17^, along-with high-throughput single-cell pancreatic islets^18–23^, have been generated. For example, Muraro et al.^23^ used the multiplexed linear amplification2 (CEL-seq2) protocol^24^ to create a human pancreatic-islet single-cell transcriptomic atlas of 2,126 cells from four donors. Xin et al.^20^ characterized 1600 cells from healthy (n=12) and T2D (n=6) donors to identify human islet cell-type specific genes. Segerstolpe et al.^21^ sequenced the transcriptomes of human islets from healthy (n=6) and T2D (n=4) donors and reported 2,209 cells. Baron et al.^25^ reported over 12,000 individual pancreatic cells from human donors (n=4) and mouse strains (n=2) which helped them uncover novel cell type-specific transcription factors, signaling receptors, and medically relevant genes. Wang et al.^26^ annotated 430 cells based on scRNA-seq on islets from healthy (n=3), child (n=2), T1D (n=1), and T2D (n=3) individuals to detect rare cellular states and population heterogeneity. However, the limited number of cells presented in the abovementioned studies in relatively small number of human donors could not provide comprehensive human pancreatic islet transcriptome data.

To address this problem, support from NIDDK enabled the Human Pancreas Analysis Program (HPAP) to continuously collect pancreatic tissues from deceased organ donors with diabetes-relevant clinical phenotypes. Data generated by HPAP continously become available through a centralized database, PANC-DB, thus enabling the diabetes research community to access these multi-dimensional data prior to any publication. HPAP investigators recently published a pancreatic islet atlas from 24 human donors consisting of three categories: healthy (n=11), AAb+ (n=8), and T1D (n=5) donors containing ~80,000 cells using single-cell transcriptomic sequencing (scRNA-seq), ~7,000,000 using cytometry by time of flight (CyTOF), and ~1,000,000 cells using imaging mass cytometry (IMC) technologies^18^. In this report, we present a new iteration of HPAP scRNA-seq data generation and analysis of ~260,000 high-quality human pancreatic islet cells from 67 donors consisting of four categories: non-diabetic (n=31), AAb+ and normoglycemic (n=10), T1D (n=9), and T2D (n=17), the largest existing scRNA-seq dataset of human pancreatic tissues. These massive and comprehensive data which can be interactively accessed on CellxGene will enable the community to advance our understanding of the molecular changes that occur in the pancreas of individuals with diabetes.

## Data Records

All pancreatic islet sequencing data have been uploaded to PANCDB^27,28^ (https://hpap.pmacs.upenn.edu/) and the interactive session can be accessed on the cellxgene database hosted on PANCDB^29^ (https://faryabi16.pmacs.upenn.edu/view/T1D_T2D_public.h5ad/). The raw islet sequencing data in the form of FASTQ files for each HPAP^28^ sample from the PANCDB database^27^. The processed data can be downloaded in two formats, RDS or h5ad, which can be loaded for analysis using Seurat^30^ and Scanpy^31^ pipelines.

HPAP^28^ is an ongoing project continuously generating new pancreatic islet tissue from human samples. As a new organ donor becomes available, each islet preparation undergoes several quality control steps, single-cell library preparation, sequencing, and the generation of FASTQ files. Finally, the raw FASTQ files will undergo the described cellranger and Seurat-based upstream and downstream pre-processing steps. An updated scRNA-seq data object will be periodically generated and automatically hosted in the PANCDB database^27^. In PANCDB, a user can obtain the expression of any gene within cell types and perform differential expression analysis between cell types on the fly using the CellxGene interactive tool.

## Technical Validation

The islets were digested into single cells within 24 to 72 hours of islet isolation from nondiabetic donors (age = 35-60 years; BMI = 25-40; A1C < 5.7), AAb+ donors (age <= 30 years), T1D donors (age <= 40; BMI <= 30), and T2D donors (age = 35-60 years; BMI 25-40; A1C > 6.5) (Fig. 1a). Detailed clinical information of donors showing their medical history, BMI, age, auto-islet antibody test, HbA1c, and C-peptide levels is provided (Supplementary Table S2). We used each individual’s medical history to determine whether the organ donor was non-diabetic, T1D, or T2D. In addition, the AAb+ screening test, including GAD, IA-2, mIAA, and ZnT8A, was performed on all donors to determine the AAb positivity. The donors with no medical history of diabetes but having AAb positivity were determined as AAb+ donors (Supplementary Table S2). We also reported quality control information about sequencing, including the number of cells, sequencing saturation, sequencing depth, and the median number of detected genes for each donor (Supplementary Table S3).

**Fig. 1.**
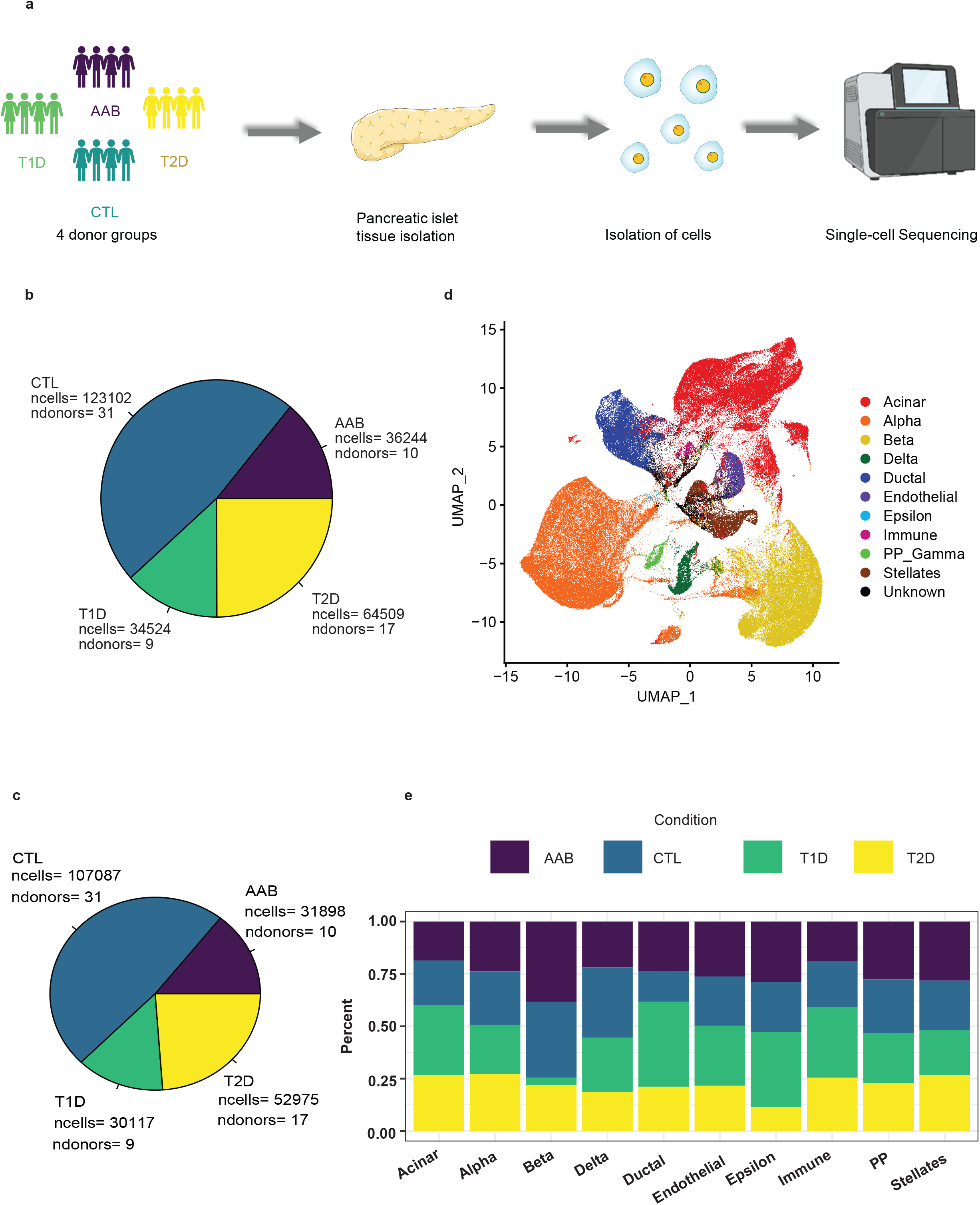
scRNA-seq reveals the cell populations of the human pancreatic islets. **a.** Overview of the scRNA-seq workflow using human pancreatic islet tissue samples. **b.** Pie chart showing the number of cells and donor distribution across different biological conditions. **c.** Uniform manifold approximation and projection (UMAP) plot showing the proper classification of islet cells. **d.** Stacked bar chart showing the percentage-wise distribution of cell types across different biological conditions.

We used cellranger to process the raw FASTQ files and generate the counts. The counts across samples were then aggregated by performing the chemistry batch correction within the cellranger pipeline. The total number of cells before pre-processing was 258,379 across 67 donors. The number of donors and cells in each condition is shown in (Fig 1.b). We used our customized Seurat pipeline for quality control (QC) and further pre-processing steps. The initial QC was performed by measuring the number of expressed genes detected, the number of UMI, and the percentage of mitochondrial genes in each cell (Fig 2.a). The number of genes detected as transcripts, UMI, and mitochondrial genes were filtered based on optimal thresholds removing the outliers, and the results are shown in Fig 2.c. The median number of genes detected as transcribed per cell across all conditions was ~2,500 (Fig 2.d). We also compared the pre-processing and median gene per cell per sample with previously published pancreatic single-cell islet data^18^ and found similar results. The percentage of mitochondrial genes observed in pancreatic islets was similar to other organs such as prostate^32^, peripheral blood mononuclear cell (PBMC)^33^, testis^34^, and liver^35^. The proportion of mitochondrial genes detected reflects the state of the cells, and inclusion and exclusion criteria are controversial. In this study, we filtered the cells with a mitochondrial gene percentage of > 25%, which is relatively conservative.

**Fig. 2.**
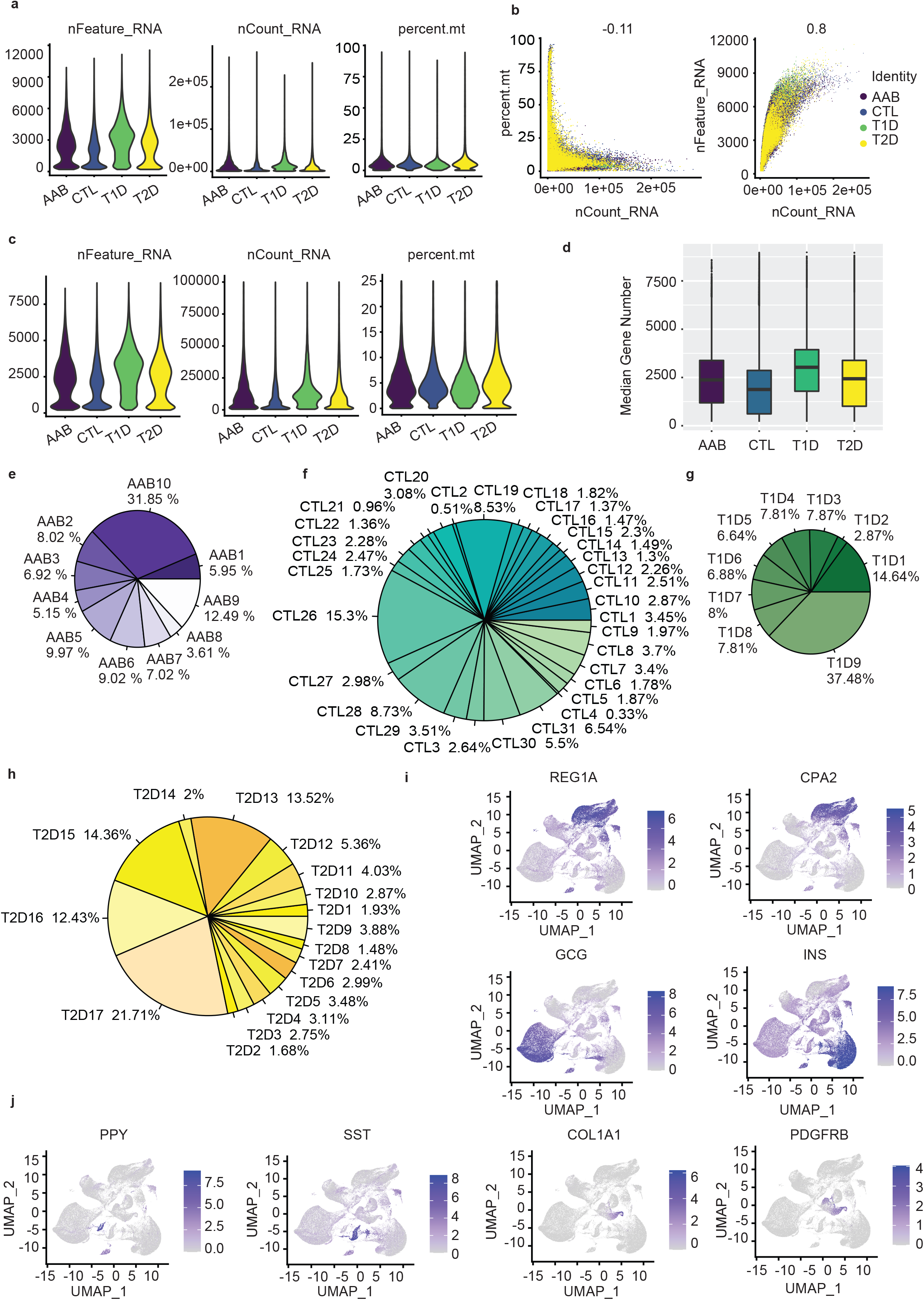
Quality control (QC) of human pancreatic islet single-cell data. **a.** Violin plots showing the number of genes, unique molecular identifiers (UMIs), and the percentage of mitochondrial genes in each cell of sixty-seven islet samples from four conditions before filtering. **b.** Scatterplots illustrate the relationship between the percentage of mitochondrial genes and the mRNA read counts and the relationship between the amount of mRNA and the read counts before filtering. **c.** Violin plots showing the number of genes, unique molecular identifiers (UMIs), and the percentage of mitochondrial genes in each cell of sixty-seven islet samples from four conditions after filtering. **d.** Median gene number showing the cell distribution across all conditions after final pre-processing. **e-h.** Pie charts showing the percentage distribution of cells across individual donors. AAB+ (e); CTL (f); T1D (g); T2D (h). **i-j.** Multiple UMAP’s depicting the validation of cell type-specific expression of marker genes.

After QC and data pre-processing, including doublet removal and normalization, the total number of cells reached to 222,077 cells across 67 donors (Fig 1.c). The percentage of cells for each donor across different conditions is shown in Fig 2.e-h. Next, based on the marker genes described in Supplementary Table S1, we utlized the scSorter method and annotated cells into 10 clusters, corresponding to acinar cells, alpha cells, beta cells, delta cells, ductal cells, endothelial cells, epsilon cells, immune cells, PP gamma cells, and stellate cells. Around 3% of cells were classified as unknown. We used the UMAP clustering approach to visualize the cell types (Fig 1.d).

Our results show that the alpha and acinar cells were abundant, with 65,113 and 59,494 cells respectively, constituting the first and second largest cell populations. The third largest cell population among the ten clusters was beta cells containing 44,559 cells. However, the percentage of beta cells in T1D donors was found to be significantly lower (Fig 1.e), as expected from the pathogenesis of T1D^18,36,37^. Our results confirm that the beta cells were indeed found to be less than < 5% in T1D, whereas they were found to be significantly higher in other conditions, such as non-diabetic controls, AAb+, and T2D (Fig 1.e). These results validate our annotation strategy of using the cell type specific marker genes and scSorter method. In addition, the cell type proportions also matched our the previously single-cell islet study^18^, where different annotation methods were applied. For additional validation, we document the expression of marker genes for the major pancreatic cell types, the acinar cells whose markers were REG1A and CPA2, and alpha cells for which the markers were GCG and INS in Fig 2.i. In addition, we also checked the expression of the markers for the less abundant cell clusters such as PP Gamma (PPY), Stellates cells (COL1A1, PDGFRB), and Delta cells (SST) (Fig 2.j). The UMAPs demonstrate well-separated cell types, validating our results. Overall, we consider our cell-type annotation results to be robust and reliable. Finally, data presented here can be interactively available through CellxGene hosted on PancDB. CellxGene exploits modern web development techniques and enables interactive visualization of large number of cells, allowing for testing hypothesis with functionalities including subsetting, differential expression, and others to help the community to explore the HPAP data. Together, our continuously expanding single-cell transcriptomic map of human pancreatic islet cells will help researchers to study islet cell biology and the relationship between cell types and disease pathologies.

## Methods

The current HPAP workflow for the analysis of pancreatic islets scRNA-seq is outlined in Fig. 1a.

### Human pancreatic islet tissue procurement and isolation

Pancreata were procured by the HPAP consortium (https://hpap.pmacs.upenn.edu/), which is part of the Human Islet Research Network (https://hirnetwork.org/) under approval from the University of Florida Institutional Review Board (IRB no. 201600029) and the United Network of Organ Sharing. Informed consent for each organ donor was obtained from a legal representative of each donor. Medical charts and C-peptide levels were measured and reviewed according to the guidelines set by American Diabetes Association. C-peptide analysis was performed as previously described^18^. AAb+ screening, including for antibodies against GAD, IA-2, mIAA, and ZnT8A, was performed on all donors before organ collection, and the AAb positivity was confirmed after tissue processing and islet isolation^18^ (Supplementary Table S2). The organs were processed and islets isolated as previously described^18,38,39^. Pancreatic islets were captured and dissociated into single cells without any specific sorting and then processed for singlecell library preparation^40^.

### 10x Genomics library preparation and cell ranger-based sample aggregation

The single-cell RNA sequencing (scRNA-seq) libraries were prepared using the Single-cell 3’ Reagent Kits v2 (10X-Chromium-GEX-3p-v2), v3 (10X-Chromium-GEX 3p-v3), and v3.1 (10X-Chromium-GEX-3p-v3.1) following the user guide (https://support.10xgenomics.com/single-cell-gene-expression/index/doc/user-guide-chromium-single-cell-3-reagent-kits-user-guide-v2-chemistry).

Illumina HiSeq 4000 or NovaSeq 6000 platforms were used for paired-end sequencing. The resulting 3’ gene expression data were processed using the 10x Genomics Cell Ranger v6.1 (CR v6.1) software^41^. The CR v6.1 comprises several pipelines that are used to process the single-cell data for aligning reads, generating feature-barcode matrices, and clustering. The main pipelines we used for processing scRNA-seq data were cellranger mkfastq, cellranger count, and cellranger aggr.

### Cellranger mkfastq, count, and aggr

The mkfastq pipeline was designed specifically for 10x Genomics libraries and depends on Illumina’s bcl2fastq2 (v 2.20) program. It takes raw base call (BCL) files generated by the sequencer and a sample sheet with Lane, Sample Id, and index for each library input. The program demultiplexes and converts the BCL files into FASTQ files. The count pipeline performs filtering, alignment (GRCh38 reference genome), barcode, and UMI counting. It can take input FASTQ files from multiple sequencing runs and uses cellular barcodes to generate feature-barcode matrices. The aggr pipeline combines count data from multiple samples of cellranger count to build an experiment-wide feature barcode matrix. By default, we performed chemistry batch correction of different libraries. The pipeline optionally offers to normalize the multiple runs to similar sequencing depth and ultimately re-computing the feature-barcode matrices. However, we performed the normalization in the downstream Seurat pipeline; therefore, the parameter was set as −none.

### scRNA-seq data pre-processing using Seurat

The merged filtered feature-barcode matrix included counts from all the HPAP samples adding up to around 260,000 cells. We built the pipeline for downstream analysis using standard scRNA-seq packages such as Seurat v4.1.0^30^, scDblFinder v1.8.0^42^, SingleCellExperiment v1.16.0^43^, sctransform v0.3.3^44^, and scSorter v0.0.2^45^ for building our pipeline carried out the following filtering steps.

### Initial Pre-processing

In the first step, we considered only those cells that contained expression values >0 for at least 200 genes using the min.features parameter. The min.cells parameter indicates to include only features detected in at least three cells.

### Doublet finding using scDblFinder

Doublets are commonly found in scRNA-seq data generated from droplet-based protocols. They arise due to cell capture or sorting mistakes, especially when thousands of cells are involved^41,43^. Doublets are undesirable in scRNA-seq data as they lead to artifactual findings. We employed the scDblFinder method to identify and remove potential doublets. The scDblFinder uses an object of class SingleCellExperiment that contains at least the counts. It trains an iterative classifier on the neighborhood of real cells and artificial doublets^42^. We measured the doublets for each donor sample separately by passing a vector of sample ids given as a parameter to the scDblFinder function. The program was run on multiple threads using the BPPARAM parameter to achieve faster computation. The scDblFinder automatically chooses the doublet rate for 10x data; therefore, we left this parameter empty. The output includes two key columns, scDblFinder.score and scDblFinder.class. While the former represents the doublet score assigned for each cell, the latter shows whether the cell was assigned as a doublet or singlet.

### Second-stage pre-processing

We further filtered the data based on nFeature_RNA and nCount_RNA, representing the number of genes detected in each cell and the total number of molecules (UMIs) detected within a cell. While the low nFeature_RNA in a cell may be dead/dying or represent an empty droplet, the high nCount_RNA and nFeature_RNA indicate that the cell may be a doublet or multiplet. This filtering, along with the filtering of mitochondrial reads (Percent.mt), is a crucial pre-processing step because removing such outliers from these groups will also remove some doublets or dead/empty droplets. In addition, we removed the cells marked as doublets from the scDblFinder in the previous step.

### Normalizing and scaling data using single cell transform (SCT)

Seurat’s SCTranform (SCT) function was used to normalize the counts that measure the differences in sequencing depth per cell for each sample. The SCT method is built on the regularized negative binomial model to perform normalization and variance stabilization of single-cell RNA-seq data^44^. It removes the variation due to sequencing depth (nUMIs). The vars.to.regress parameter allowed us to regress the variation from other sources, such as the percentage of mitochondrial reads. The output of the SCT model is the normalized and scaled expression levels for all the transcripts. Lastly, the variable.features.n parameter is used to select the variable features and is set as 3,000.

### PC calculation and clustering

We performed principal-component analysis (PCA) based on the top 3,000 most variable genes obtained from the SCT transformed data. We first calculated 50 PCs, and then the dimensionality was determined using an elbow plot showing each PC’s standard deviation. Based on the elbow plot, we chose the top 20 PCs.

The FindNeighbors function in Seurat was used to compute the k-nearest neighbors in the dataset. The calculations were made on the reduced dimensions of the 20 PCs determined based on the elbow plot in the previous step. After calculating k-nearest neighbors, the FindClusters function. built on the concept of shared nearest neighbors (SNN), clustering algorithm was used to cluster the cells at the resolution of 0.8. Finally, the uniform manifold approximation and projection (UMAP) dimensional reduction technique was run on the selected 20 PCs.

### Automatic cell annotation using the single-cell sorter method (scSorter)

We employed the marker-based scSorter cell annotation method to assign cells to known cell types based on the marker genes. The cell type annotations were made based on the marker genes reported in previous studies^4,18,20,21,23,26^ (Supplementary Table S1). The scSorter method is based on a semi-supervised learning algorithm. It does not require all genes but only a set of marker genes and highly variable genes from the expression data to perform the analysis^45^. The scSorter function expects the preprocessed data as input. Therefore, we used the previously normalized, scaled, and transformed expression matrix generated by the SCT method as input. The top 3,000 most variable genes selected by the SCT method were also given as input for the scSorter method. Additionally, the scSorter function accepts “weight” for the marker genes. Therefore, we assigned equal weights to all markers through the default_weight option.

## Code Availability

The scripts used for data processing and analysis of the scRNA-seq data is available on GitHub^46^ https://github.com/faryabiLab/HPAP-scRNA-seq-Workflow-2022. Cellxgene interactive tool with HPAP data can be accessed cellxgene database^29^ (https://faryabi16.pmacs.upenn.edu/view/T1D_T2D_public.h5ad/).

## Acknowledgements

We thank helpful discussions from Vahedi, Faryabi, and Kaestner lab members. This work was supported by National Institute of Health grants UC4 DK112217 and U01DK112217, R01HL145754, U01DK127768, U01DA052715, the Burroughs Wellcome Fund, the Chan Zuckerberg Initiative, W. W. Smith Charitable Trust, the Penn Epigenetics Institute, and the Sloan Foundation awards to G.V.

## Author contributions

A.R.P performed single-cell RNA-seq analysis, made figures, and wrote the original draft; G.V conceived and supervised the project; K.K supervised single-cell RNA-seq data generation. J. S., R.B. F and G.V. supervised data analysis. A.R.P and G.V reviewed and edited the manuscript. All authors have read and agreed to the published version of the manuscript.

## Additional information

**Correspondence** and requests for materials should be addressed to G.V

## Table Legends

**Supplementary Table S1.** Pancreatic islet cell type markers were used for annotations.

**Supplementary Table S2.** Clinical information of all HPAP organ donors.

**Supplementary Table S3**. Summary sequencing statistics of all sixty-seven HPAP samples.

